# Rv0687 a Putative Short-Chain Dehydrogenase is indispensable for pathogenesis of *Mycobacterium tuberculosis*

**DOI:** 10.1101/2023.12.12.571312

**Authors:** Gunapati Bhargavi, Mohan Krishna Mallakuntla, Deepa Kale, Sangeeta Tiwari

## Abstract

*Mycobacterium tuberculosis (Mtb),* a successful human pathogen, resides in host sentinel cells and combats the stressful intracellular environment induced by reactive oxygen and nitrogen species during infection. *Mtb* employs several evasion mechanisms in the face of the host as a survival strategy, including detoxifying enzymes as short-chain dehydrogenases/ reductases (SDRs) to withstand host-generated insults. In this study, using specialized transduction we have generated a Rv0687 deletion mutant and its complemented strain and investigated the functional role of Rv0687, a member of SDRs family genes in *Mtb* pathogenesis. Wildtype (WT) and mutant *Mtb* strain lacking Rv0687 (RvΔ0687) were tested for *in-vitro* stress response and *in-vivo* survival in macrophages and mice models of infection. The study demonstrates that Rv0687 is crucial for sustaining bacterial growth in nutrition-limited conditions. The deletion of Rv0687 elevated the sensitivity of *Mtb* to oxidative and nitrosative stress-inducing agents. Furthermore, the lack of Rv0687 compromised the survival of *Mtb* in primary bone marrow macrophages and led to an increase in the levels of the secreted proinflammatory cytokines TNF-α, and MIP-1α. Interestingly, the growth of WT and RvΔ0687 was similar in the lungs of infected immunocompromised mice however, a significant reduction in RvΔ0687 growth was observed in the spleen of immunocompromised Rag^-/-^ mice at 4 weeks post-infection. Moreover Rag^-/-^ mice infected with RvΔ0687 survived longer compared to WT *Mtb* strain. Additionally, we observed significant reduction in bacterial burden in spleens and lungs of immunocompetent C57BL/6 mice infected with RvΔ0687 compared to complemented and WT *Mtb* strains. Collectively, this study reveals that Rv0687 plays a role in *Mtb* pathogenesis.

## 1. Introduction

*Mycobacterium tuberculosis* (*Mtb*) is the causative agent of tuberculosis (TB), a bacterial disease that claims about 1.5 million lives worldwide in 2022 (1). On infection, *Mtb* is phagocytosed by sentinel cells of the host, macrophages. These cells are equipped with cell autonomous innate immune responses, which act as front-line defenders against *Mtb* infection (2,3). Activated macrophages upregulate various antimicrobial pathways including the expression of inducible nitric oxide synthase (INOS), phagocyte NADPH oxidase (NOX2/gp91^phox^) and xanthine oxidase system that gets recruited to the *Mtb* phagosome and generates antimicrobial reactive nitrogen and oxygen species (ROS and RNS) vis respiratory burst to kill *Mtb* (3–6). Mice deficient in both *phox* and *iNOS* are highly susceptible to *Mtb* infection that shows RNS and ROS generation is required by the host for protection against *Mtb* (7). These molecules alter the intracellular signaling events, triggering the host antimicrobial responses that primarily aim to eliminate *Mtb* (2,8). The host cell ROS and RNS interacts directly with its bacterial targets as well during the process get converted into chemically district oxidants as hydrogen peroxide (H_2_O_2_), hypochlorites (HClO), peroxynitrite (ONOO^-^) and hydroxy radicals (9,10). These reactive species damage *Mtb* DNA, oxidize lipids and proteins, including highly sensitive proteins as iron-sulfur (4Fe-4S) cluster proteins (9). Thus, *Mtb* infection prompts macrophages to generate ROS and RNS as a mechanism to kill intracellular bacteria (10,11). The ROS and RNS production by *Mtb-*infected macrophages, as part of host cell-autonomous innate immune response, not only play a role in direct host defense against the bacteria but also trigger signaling pathways that contribute to the overall antimicrobial responses of these immune cells (12).

Survival of *Mtb* in the host is does not only depends on the manipulation of the host but also on its ability to encounter stresses imposed by immunologically active macrophages. *Mtb* employs several detoxifying antioxidant thiols and enzyme systems, such as superoxide dismutase (SOD), catalase-peroxidase, alkylhydroperoxidase (AhpC), and thioredoxin reductase (9,13). However, the gene encoding global stress regulator *oxyR*, commonly found in several other bacteria, is absent in *Mtb*. The absence of *oxyR* in *Mtb* is suggested to be compensated by the presence of several AhpC, which help the bacteria tolerate ROS and RNS induced by the infected host cells (14,15). In addition to AhpC, recent research has revealed that *Mtb* possesses a category of enzymes known as short-chain dehydrogenase/reductases (SDRs), which aid in managing peroxidase stress, detoxification, *in-vitro*, *in-vivo* survival, drug resistance and overall homeostasis (16,17). The SDRs are one of the largest protein families, distributed widely in several organisms and classified into seven families such as classical, atypical, extended, intermediate, divergent, complex, and unassigned (18,19). SDRs can utilize a range of diverse substrates, such as alcohols, sugars, and aromatic compounds (19). SDRs represent various enzyme classes such as isomerases, lyases and oxidoreductases, although the majority of the SDRs are oxidoreductases (20,21). These SDRs share an NAD(P)^+^ Rossmann fold domain for substrate binding and use NADH or NADPH as its electron donors (21). Moreover, SDRs are involved in metabolic pathways such as detoxification, drug resistance, steroid metabolism, and biosynthetic pathways of bacteria (22,23).

In the context of *Mtb* infection, specific SDRs have been implicated in modulating the activation or inactivation of certain drugs used in TB treatment, highlighting the involvement of SDRs in bacterial physiology and survival (24,25). The genome of *Mtb* possess 11 SDRs, namely Rv3057c, Rv0851c, Rv1856c, Rv3485c, Rv3530c, Rv3548c, Rv3559c, Rv2509, Rv3224, Rv0148, Rv1484 and Rv0687 (as listed in Tuberculist) (26). Rv0687 has been shown to have homologs in other mycobacteria that are characterized as mycofactocin-associated dehydrogenases with non-exchangeable NAD-cofactors and as a putative carveol dehydrogenase (27). Furthermore, the Rv0687 transposon mutant is attenuated in both mouse and primate designer assays for defined mutant analysis (28).

Interestingly, in our RNA-seq studies (data not shown) we found that Rv0687 is highly upregulated in *Mtb* treated with NMR711, a compound that we found can inhibit the growth of both drug-sensitive and multidrug-resistant TB (29). However, precise knockout deletion mutant of Rv0687 is not generated and its role in *Mtb* pathogenesis and antibiotic resistance is under investigated. Therefore, this study involves characterizing the role of Rv0687 in the pathogenesis of the *Mtb*. Notably, Rv0687 is a non-essential in the *Mtb* genome, with a gene length of 828 base pairs, encoding a 275 amino acid protein with a molecular mass of 28kDa. In the current study, we generated a knockout mutant of Rv0687 (RvΔ0687) using specialized transduction and characterized the phenotype of the mutant in comparison with the wild type (WT) and complemented *Mtb* strains (C-Rv0687). We demonstrated that Rv0687 is required by *Mtb in-vitro* to cope up with the stresses as nutrient-limiting conditions, oxidative and nitosative stress, as well as for survival and growth in mouse bone marrow-derived macrophage (BMDM) infection assays. Moreover, it is required by *Mtb* for survival and proliferation *in-vivo* in immunocompetent and immunocompromised mice models of infection.

## 2. Materials and methods

### 2.1. Bacterial strains

Wild type *Mtb* bacterial strains and vector plasmids (p004s, phAE159, pMV361) used in this study were gift from Dr. William R Jacob Jr (Albert Einstein College of Medicine) and are listed in Supplementary materials. All the Mtb strains were grown at 37 ^ο^ C in Middlebrook 7H9 (Difco) media, supplemented with 10% OADC (oleic acid-albumin-dextrose-catalase), 0.5% glycerol, and 0.05% tyloxapol. To test the growth kinetics, *Mtb* strains were grown in 7H9-Dextrose (minimal media) with 0.5% glycerol and tyloxapol. To grow *Mtb* strains carrying antibiotic markers, hygromycin (75μg/mL) or kanamycin (20μg/mL) were supplemented to the media wherever needed.

### 2.2. Gene knockout construction and confirmation

Gene knockout of *Mtb* was performed by specialized transduction as described previously (30). Briefly, the flanking regions of the Rv0687 gene sequence were obtained from Tuberculist and cloned into the p0004-SacB vector, carrying the Hyg^R^ resistance gene for homologous recombination. The recombinant construct was further cloned into phAE159 temperature-sensitive phasmid and a knockout mutant of Rv0687 was generated using specialized transduction. Deletion of Rv0687 was confirmed using 3-primer strategy (31) (Fig. S1A), and whole genome sequencing performed on a MiSeq instrument (Illumina) following the protocol provided by Illumina. Further, the mutant strain was complemented by integrating the wild-type copy of the gene into the genome of the RvΔ0687 mutant using a pMV361 mycobacterial integrative vector and complementation was confirmed gene-specific primers (**Fig.S1B**) and kanamycin primers (31). The details of the strains, plasmids, phasmids, and primers used in the study are provided in supplementary tables (**Table S1-S3**).

### 2.3. Assessing *Mtb* response to ROS and RNS generating agents

The wild-type, RvΔ0687 and C-Rv0687 *Mtb* strains were grown to OD_600 nm_ ≍ 0.8-1 and diluted to 0.05×10^^^7 cells/mL with 7H9 dextrose media to a final volume of 5mL. Diluted bacterial cultures were treated with ROS; 5 or 10mM hydrogen peroxide (H_2_O_2_) and 2mM tert-butyl hydroperoxide (tBOOH) or RNS; 2mM S-nitroso-N-acetyl-D, L-penicillamine (SNAP) and 2 and 10mM sodium nitrite (SN) generating agents; and the cultures were incubated for 24, 48, 72 and 96 h after post-treatment and the colony forming units (CFUs) were determined by plating the serially diluted bacterial culture onto 7H10 plates and incubating the plates at 37 ^ο^C for 4-6 weeks.

### 2.4. Quantification of *Mtb* NADH and NAD^+^ levels

NADH-binding proteins were extracted using the NAD/NADH assay kit from Abcam. Briefly, 0.8 to 1 OD_600_ of wild type, RvΔ0687 and C-Rv0687 strains were lysed using NAD lysis buffer, and the resulting proteins were quantified by adding an NADH developer that reacted with the proteins to produce a calorimetric signal that was measured at 450 nm using a spectrophotometer (Spectramax). The concentrations of NADH/NAD**^+^** were estimated from the resulting spectra, allowing for the calculation of the NADH/NAD**^+^** ratio (32).

### 2.5. Minimal inhibitory concentration determination by alamar blue assay

The wild type, RvΔ0687 and C-Rv0687 bacterial strains were grown in 7H9 complete media to OD_600 nm_ ≍ 0.5-0.8. To determine the minimal inhibitory concentration (MIC) of Isoniazid (4-0.03μg/ml), Delamanid (4-0.03 μg/ml), NMR711 (200-1.5 μg/ml) and Bedaquiline (2-0.015 μg/ml), the drugs were serially diluted 2-fold to reach the least concentration gradient. 100μl of bacteria at an OD_600 nm_ of 0.1 was added into the wells containing the drugs and followed by the addition of alamar blue and incubated the plates at 37 ^ο^ C for 24, 48, 72 and 96h. At the indicated time points plates were assessed for the bacterial growth and were read in a spectrophotometer at a wavelength of 560/590nm.The MIC is assessed based on color change of alamar blue.

### 2.6. Infection of bone marrow derived macrophages (BMDMs)

The C57BL6 mice were euthanized, and the femurs and tibias were separated and flushed thoroughly with Dulbecco’s modified Eagle Medium (DMEM; Invitrogen) to obtain cells. After lysing of red blood cells, the cells were maintained in DMEM, containing 10% FBS and 30% L929 supernatant (complete media) (33). Cells were cultured in tissue culture flasks for 7 days and seeded at 0.25 x10^6^ cells per well into 24-well plates. The bacterial strains were grown at 37 ^ο^ C to OD_600 nm_ ≍ 0.8-1. After washing with phosphate-buffered saline with tyloxapol (PBST), the bacterial suspensions were diluted in DMEM and used to infect BMDMs cells for 3h at 37 ^ο^ C in 5% CO_2_ at a multiplicity of infection of 5 (MOI 1:5). Following infection at 3h, the media were removed, and the wells were washed with PBST before treating with gentamycin for 1h. Subsequently, the antibiotic media was removed, and the wells were washed with PBST and replenished with fresh DMEM media (complete media). At every time point tested, the infected BMDMs were lysed with Triton X-100, and the lysates were serially diluted in PBST and plated on 7H10 agar media to determine the number of colony-forming units (CFUs). The post-infected cell supernatants were collected before lysis of the cells and analyzed for IP-10, IL-1β, MIP-1 and TNF-α cytokines.

### 2.7. Measurement of ROS and RNS in BMDMs during infection

To assess the levels of ROS generated during BMDM infection, 0.1% Nitroblue Tetrazolium Chloride (NBT; Sigma) is added to wild type, RvΔ0687 and C-Rv0687 infected BMDMs in the 96 well plate and incubated at 37 ^ο^ C for 30 min. After incubation, the NBT solution was removed, and the cells were suspended in 2M potassium hydroxide to stabilize the color. Dimethyl Sulfoxide (DMSO) was then added to solubilize the formazan. The amount of formazan, which represents the ROS levels, was quantified by measuring the optical density at 570 nm against DMSO as the reference.

To assess the levels of RNS, post-infected supernatants from wild-type, RvΔ0687 and C-Rv0687 infected BMDMs were collected, and the assay was performed using the Griess Reagent Kit (Biotium) following the guidelines as provided in the kit. Various concentrations of sodium nitrite references were prepared to create a standard curve, which was used to measure the absorbance at 570 nm. Post-infected supernatants were mixed with deionized water and Griess reagent, and the reaction mixture was incubated for ≍30 minutes at room temperature. The absorbance was then measured at 570 nm, and the optical density reading represented the nitrite concentration in the samples. These values were normalized to the reference values.

### 2.8. Mouse infection experiments

The wild type, RvΔ0687 and C-Rv0687 strains were cultured to an OD_600 nm_ ≍ 0.8-1. The bacteria were washed twice with PBST. Subsequently, the samples were sonicated in a Branson cup-horn sonicator, twice for 10 seconds each time, and then diluted to achieve the desired cell densities of ∼1×10^6^ CFUs/ml. C57BL/6 mice (6-8 weeks old) were obtained from the Jackson Laboratory, and Rag^-/-^ mice (6-8 weeks old) were obtained from the in-house breeding facility. The mice were infected with a dose of *Mtb* strains (200-250 CFUs per lung) using a Glass-Col chamber as described previously (34) and were euthanized at indicated time points. Lung and spleen were harvested and homogenized with PBST. The CFUs were performed by serially diluting the lysate and plated onto 7H10 plates and colonies were enumerated after 4-6 weeks of incubation at 37 ^ο^ C and CFUs were estimated per lung and spleen. The animal protocol was approved by the University of Texas at El Paso Institutional Animal Care and Use Committee (IACUC) protocol, A-202004-3 as per NIH principles for animal usage.

## 3. Results

### 3.1. Growth kinetics of RvΔ0687 in Broth media

To assess the growth kinetics of WT, RvΔ0687 and C-Rv0687, strains were grown in both 7H9 complete and 7H9 dextrose media, and their growth was monitored by CFU assay for 12 days. In 7H9 complete media, the growth of RvΔ0687 showed no significant differences compared to the WT and C-Rv0687 at any of the tested time points (**Fig. 1A**). Whereas, in 7H9-dextrose media, all the strains were able to survive without any change in growth till day 4, followed by a gradual decline in CFUs which is not statistically significant at day 8. However, no CFUs were obtained for the RvΔ0687 at day 12, whereas WT and C-Rv0687 were able to persist in the 7H9 dextrose media and form CFUs (**Fig. 1B**). To conclude, the failure of the RvΔ0687 growth might be due to the absence of essential nutrients for bacterial survival. The minimal media showed the difference in growth response among the *Mtb* strains.

**Figure 1.**
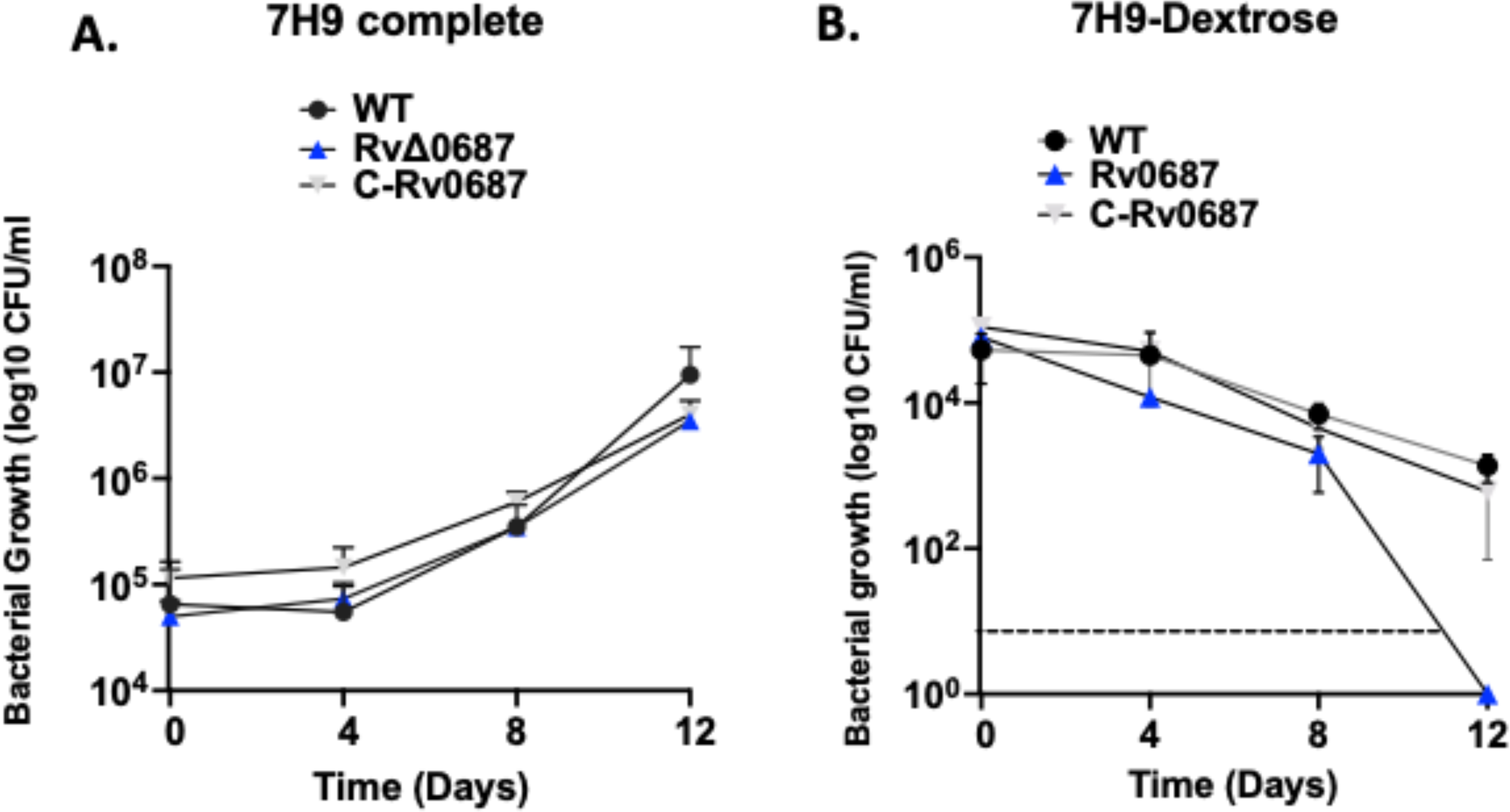
(A-B) (A) Growth kinetics in 7H9 complete. The WT, RvΔ0687 and C-Rv0687 strains were grown in 7H9 complete media, and the growth kinetics was measured at Day 0, 4, 8 and 12. Aliquots of bacterial cultures were serially diluted and spotted onto 7H10 complete media and colony forming units (CFUs) were calculated per ml of bacterial culture and graphs were plotted. (**B) Rv**Δ**0687 has growth defect in 7H9 minimal media.** The WT, RvΔ0687 and C-Rv0687 strains were grown in 7H9 dextrose minimal media till Day 12, and the growth was observed by serially diluting the bacterial strains and plating onto 7H10-OADC media. The observed colonies were enumerated to calculate CFUs/ml and graphs were plotted. Experiments were performed in triplicates, mean±SD is plotted, and significance was calculated using Two-way ANOVA between the bacterial strains at tested time points.

### 3.2. RvΔ0687 is sensitive to ROS and RNS generating agents

The response of WT, RvΔ0687 and C-Rv0687 to *in-vitro* ROS was tested using 5mM and 10 mM H_2_O_2_ and 2mM tBOOH. The RvΔ0687 showed sensitivity to exposure to 5mM and 10mM H_2_O_2_ as determined by the reduction in CFUs, compared to the WT and C-Rv0687. However, at 5mM H_2_O_2_ exposure, a significant reduction in RvΔ0687 CFUs was observed only at 96h, whereas at 10mM H_2_O_2_ exposure, at 24h decrease in CFUs was observed but at 48h the CFUs of RvΔ0687 dropped by 2 logs compared to WT and C-Rv0687 strains. Furthermore, RvΔ0687 was so susceptible to H_2_O_2_ stress that we recovered no CFUs at 96h post-treatment compared to the WT and C-Rv0687 **(Fig.2 A and B)**. In addition, the RvΔ0687 was sensitive to 2mM tBOOH at 24h, with significantly declined CFUs compared to WT and the C-Rv0687 strains. However, none of the strains were able to sustain their growth after 48h, as observed by the lack of CFUs, in the presence of 2mM tBOOH (**Fig. 2C)**. The RNS response was assessed using 2mM SNAP and 2 or 10mM SN. At 72 and 96h post-treatment, we observed a log decrease in the RvΔ0687 compared to WT and the C-Rv0687 strains in response to 2mM SNAP treatment (**Fig. 2D)**. However, neither of the tested bacterial strains (WT, RvΔ0687 and C-Rv0687) showed any difference in CFU on treatment with 2 or 10mM SN **(Fig. S2)** or in response to metal ions (copper, zinc, and iron) stress at any of the tested time points after post-treatment (**Fig. S3)**.

**Figure 2.**
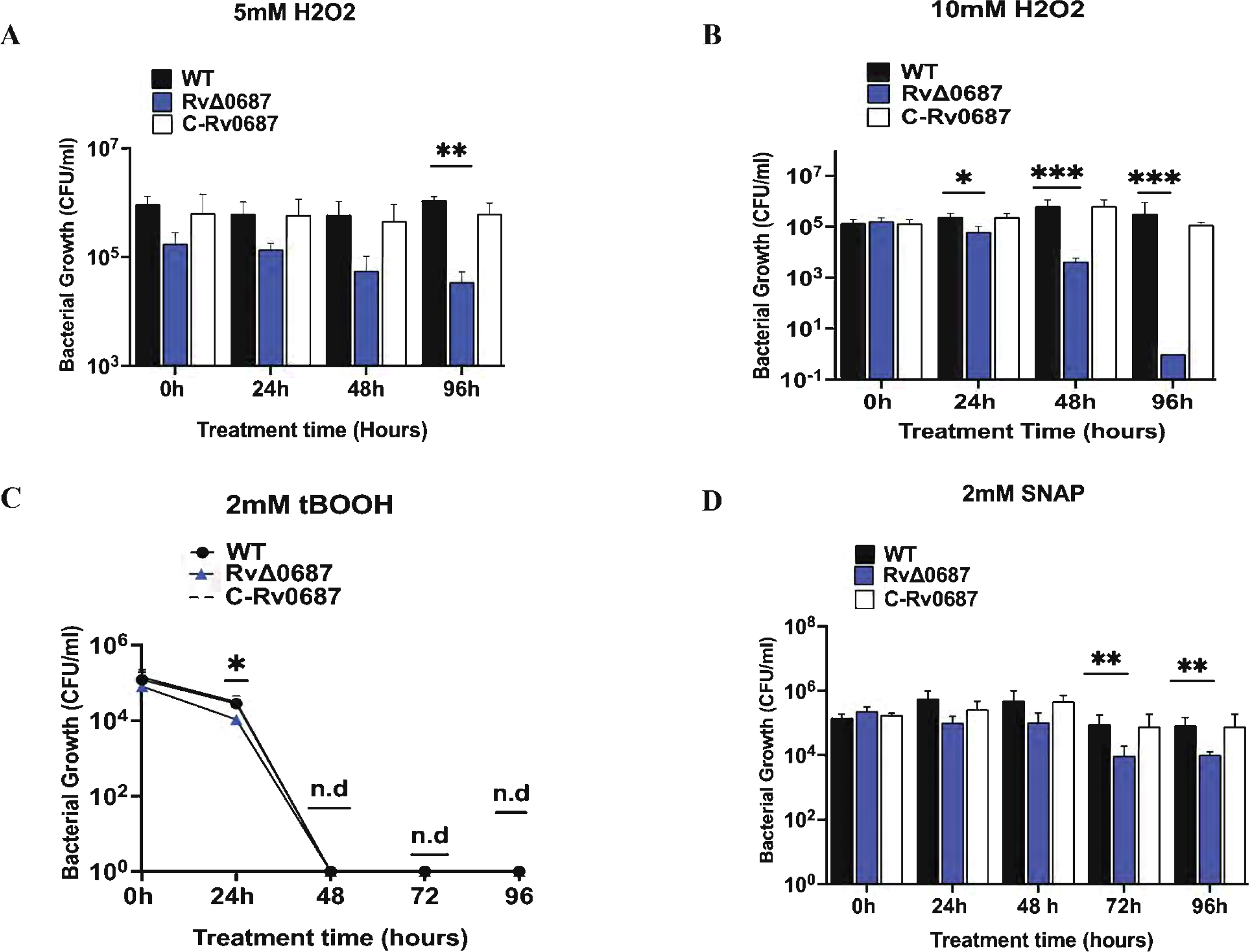
(A-D) RvΔ0687 is susceptible to oxidative (ROS) and nitrosative (RNS) stress. **(A) Effect of 5mM H_2_O_2_ stress.** The WT, RvΔ0687 and C-Rv0687 strains were grown in 7H9 complete media, pelleted, washed with PBST, and added to 7H9 dextrose media and exposed to 5mM H_2_O_2_ and the growth of all the strains were monitored at 0, 24, 48 and 96h post-treatment. The cultures were serially diluted at each time point after post-treatment and spotted onto 7H10 OADC plates and CFUs were calculated for 1ml of bacterial culture. (**B) Effect of 10mM H_2_O_2_ stress.** The WT, RvΔ0687 and C-Rv0687 strains were grown in 7H9 dextrose media and exposed to 10mM H_2_O_2_ and the growth of all the bacterial strains was monitored. The experiment was carried out in triplicates and significance was calculated using Two-way ANOVA, the error bars indicate the standard deviation (\**P<0.1* \*\**P<0.01*\*\*\**P<0.001)*. (**C) Effect of 2mM tert-butyl hydroxide (tBOOH).** The WT, RvΔ0687 and C-Rv0687 strains were grown in 7H9 complete media. The pellet was washed twice with PBST and suspended in 7H9 dextrose media, and treated with 2mM tBOOH and monitored at 0, 24, 48, 72 and 96h after post-treatment. The sensitivity was analyzed by plating the serially diluted cultures in PBST and spotted onto 7H10. CFUs were enumerated per ml of bacteria and graphs were plotted. The experiment was carried out in triplicates and significance was calculated using Two-way ANOVA and T-test between bacterial strains (\**P<0.1)*. (**D) Effect of Reactive Nitrogen Stress (2mM SNAP).** The WT, RvΔ0687 and C-Rv0687 strains were grown in 7H9 dextrose media and treated with 2mM SNAP, and the bacterial strains were spotted onto 7H10 OADC by serially diluting the cultures. The CFUs were estimated after 3 weeks of incubation and CFUs were calculated per ml of bacterial culture. The experiment was carried out in triplicates and using Two-way ANOVA significance was calculated significance between the bacterial strains (\*\**P<0.01)*.

### 3.3. Deletion of Rv0687 gene does not impact the Mtb NADH, and NAD^+^ and NADH/NAD ratio *in-vitro*

The NADH and NAD^+^ ratios have a crucial role in the redox homeostasis of a bacterial cell. Consequently, we measured the concentrations of NADH (**Fig.3 A)** and NAD^+^ (**Fig.3 B)**as well as the NADH/NAD^+^ ratio (**Fig.3 C)** for the WT, RvΔ0687 and C-Rv0687 strains. Based on our observations, it is evident that the deletion of Rv0687 has no significant impact on modulating the NADH or NAD^+^ levels and redox homeostasis of *Mtb* in aerobic conditions.

**Figure 3.**
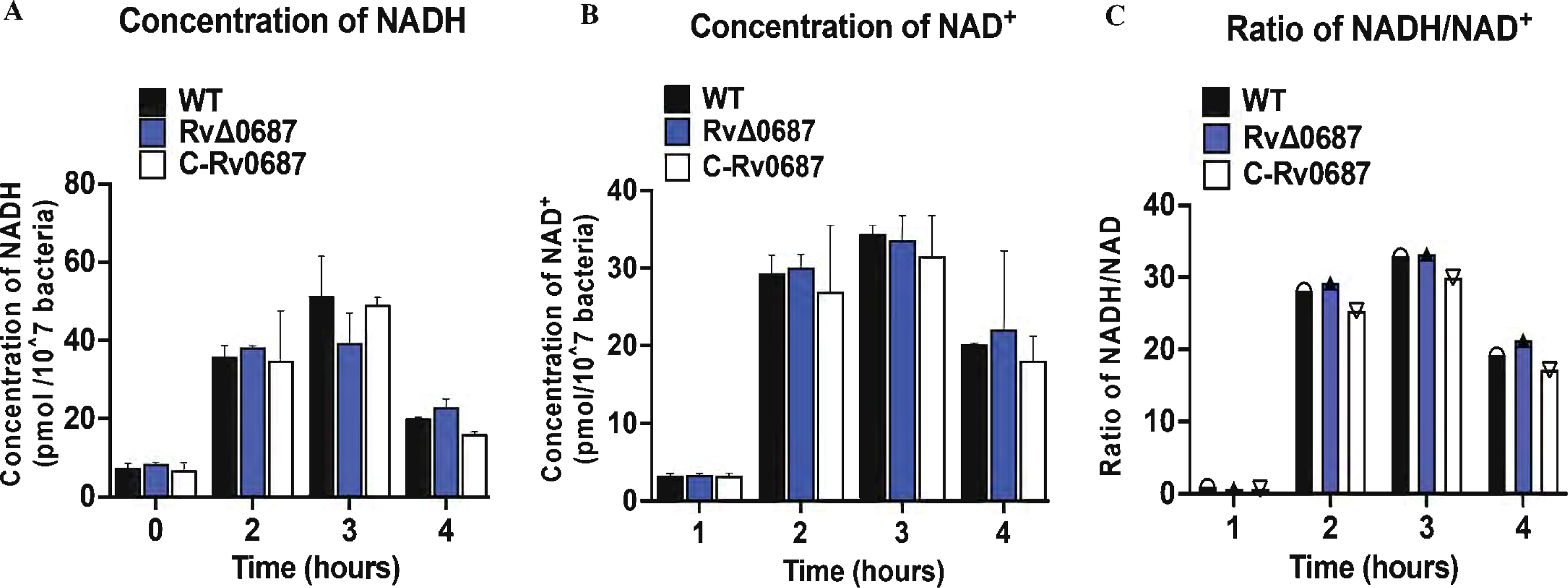
(A-C) Rv0687 deletion in *Mtb* does not impact intracellular NADH/NAD^+^ concentrations The WT, RvΔ0687 and C-Rv0687 were grown to 10^7^ and the cells were pelleted and lysed. NADH/NAD^+^ were extracted and the concentrations were estimated individually and then the ratios of NADH/NAD^+^ were calculated using T-test.

### 3.4. Deletion of Rv0687 enhances susceptibility of *Mtb* to antimycobacterial delamanid and NMR711

To assess the sensitivity of WT, RvΔ0687 mutant and C-Rv0687 strains to Isoniazid (INH), Delamanid (DEL), NMR711, Bedaquiline (BDA), and Rifampicin (RIF), and Alamar blue assay was performed. The RvΔ0687 displayed increased susceptibility to NMR711 (MIC >0.390 μg/ml) which is approximately 128-fold higher than the WT and the C-Rv0687 strains (>50 μg/ml). Interestingly, RvΔ0687 also displayed MIC of (>0.062 μg/ml) for delamanid, a recently approved TB drug, which is 16-fold higher than WT and the C-Rv0687 strains (>1 μg/ml) (**Table-1** and **Fig. S4**). Conversely, RvΔ0687 showed similar susceptibility to antimycobacterial INH, BDA, and RIF; compared to the WT and C-Rv0687 strains.

### 3.5. RvΔ0687 is required for intracellular replication and suppression of early inflammatory immune response in macrophages

The *in-vitro* survival of WT, RvΔ0687 and C-Rv0687 strains were tested using mouse BMDMs. As shown in (**Fig. 4)** no significant difference is observed in the uptake of the WT, RvΔ0687 and C-Rv0687 strains by BMDMs. Interestingly at day 1 though we observed small differences in the CFUs between WT and RvΔ0687 strains, but the differences were significant at later stages at day 3, 5 and 7 post infections (**Fig. 4 A)**. We found that Rv0687 is required for the intracellular survival of *Mtb*, as the deletion mutant of Rv0687 is attenuated for survival in the macrophages. We further assessed the production of RNS (Griess assay) and ROS (NBT assay) by BMDMs in response to infection with WT, RvΔ0687, and C-Rv0687 strains. Though there was a significant change in RNS and ROS production by WT infected vs uninfected BMDMs, but we didn’t observe any significant change in the production of ROS or RNS by BMDMs on infection with either WT, RvΔ0687, and C-Rv0687 strains (**Fig. S5 A-B)**.

**Figure 4.**
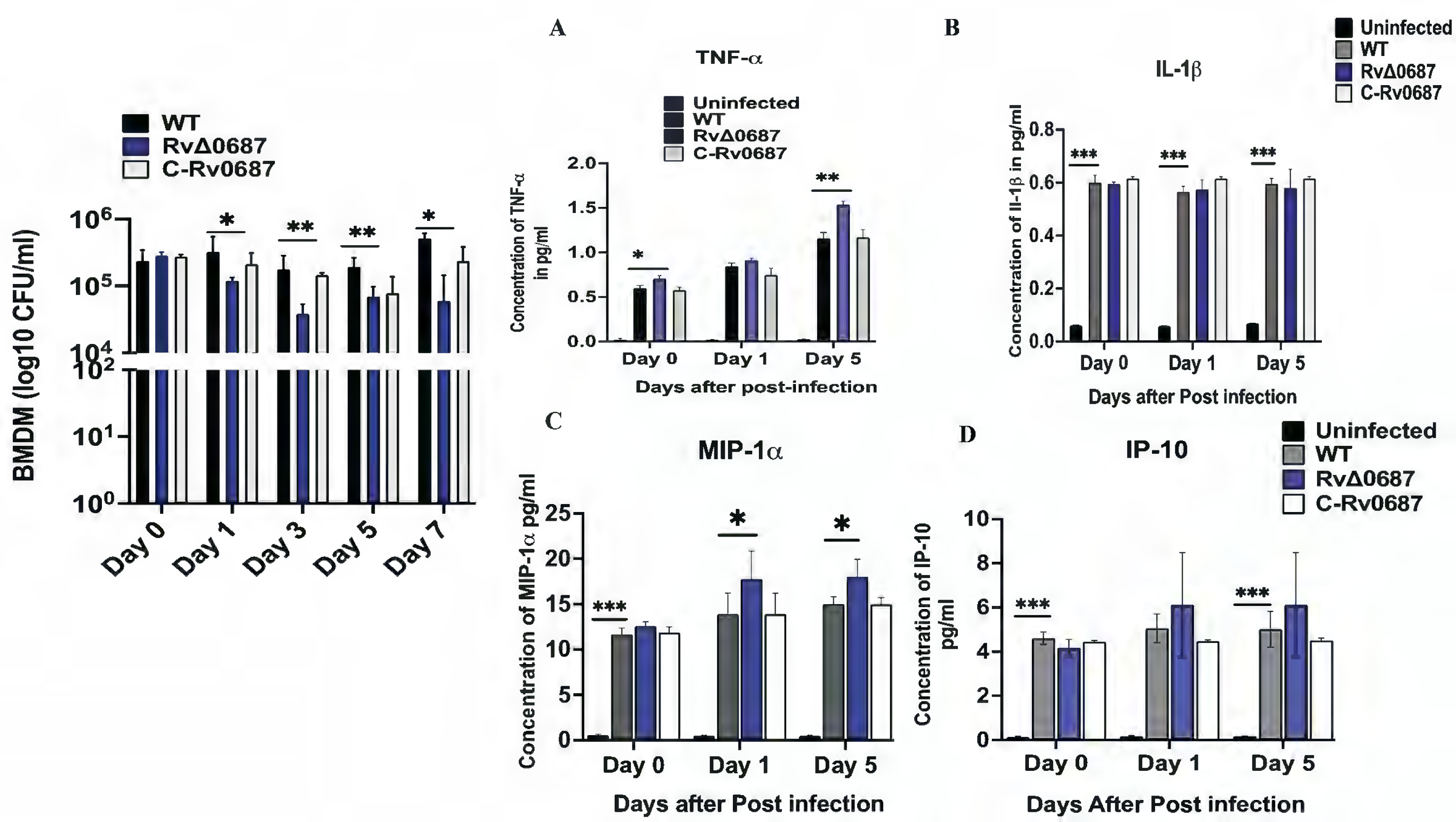
Rv0687 is required for *Mtb* growth and suppression of inflammatory immune response in BMDMs. The WT, RvΔ0687 and C-Rv0687 strains were used for BMDMs infection at an MOI 1: 5. The *in-vitro* survival was assessed after days 0, 1, 3, 5 and 7 post-infection and bacterial growth was assessed by serially diluting the infected lysate and plated on 7H10 agar and the plates were incubated for 4-6 weeks to calculate the CFUs. The experiment was performed in triplicates and Two-way ANOVA was used to calculate the significance between the bacterial strains. Figure 4**. (A-D). Cytokine Expression during BMDMs infection:** The WT, RvΔ0687 and C-Rv0687 strains infected with BMDMs were analyzed for the expression of TNF-α, IL-1β, MIP-1α, and IP-10 cytokines using lysates at Day 0, 1 and 5 post-infections. (**A)** TNF-α, (**B)** IL-1β, (**C)** MIP-1α, (**D)** IP-10. The experiment was performed in triplicates and Two-way ANOVA with the Tukey correction method is used to calculate the significance between uninfected control and infected bacterial strains (\*\*\**P<0.001* and \*\**P<0.01)*.

Further we measured the cytokines to determine the activation status of the macrophages. The expression levels of TNF-α, IL-1β, IP-10 and MIP-1α cytokines were analyzed in the WT, RvΔ0687, and C-Rv0687 strains in the cell-free post-infected supernatants of infected BMDMs at 1-5 days of post infection. Within 3h of infection compared to uninfected, WT-infected BMDMs showed 5-fold increased secretion of TNF-α, IL-1β and ∼10-fold increase in MIP-1α and IP-10.

Interestingly, RvΔ0687 infected BMDMs showed a significantly higher expression of TNF-α and MIP-1α at d1 and d5 days compared to WT and C-Rv0687 infected BMDMs (**Fig. 4 A and C**). We have not observed any significant change in the secretion of IL-1β and IP-10 by RvΔ0687 infected BMDMs compared to that of WT and C-Rv0687 infected BMDMs (**Fig.4 B and D)**.

### 3.6. RvΔ0687 is attenuated *in-vitro* in immunocompromised and immunocompetent mice

The *in-vivo* survival of WT, RvΔ0687 and C-Rv0687 strains were analyzed in immunocompromised (Rag^-/-^) and immunocompetent (C57BL/6) mice **(Fig. 5A).** Immunocompromised Rag^-/-^ mice were infected via aerosol with a dose of 200-250 CFU/mice. Mice were sacrificed at day 0, W4 and W8 to determine the CFU burden in the lungs and spleen. A group of mice infected with each strain were kept for survival studies. At W4 and W8 post-infection, WT, RvΔ0687 and C-Rv0687 showed similar bacterial burden in the lungs of Rag-/- mice **(Fig. 5B),** but in spleen at W4 the growth of RvΔ0687 was 8-fold lower than WT and C-Rv0687 strains **(Fig. 5C).** Furthermore, we compared the survival of Rag^-/-^ mice after infection with WT, C-Rv0687 or RvΔ0687 strains. Interestingly, the WT or C-Rv0687 infected group started to die between 60-70 days post-infection, while the RvΔ0687 infected animals survived till 125 days post-infection (**Fig.5D**). This suggests that RvΔ0687 is impaired in virulence and plays a significant role in the *in-vivo* survival of *Mtb* in Rag^-/-^ mice.

**Figure 5.**
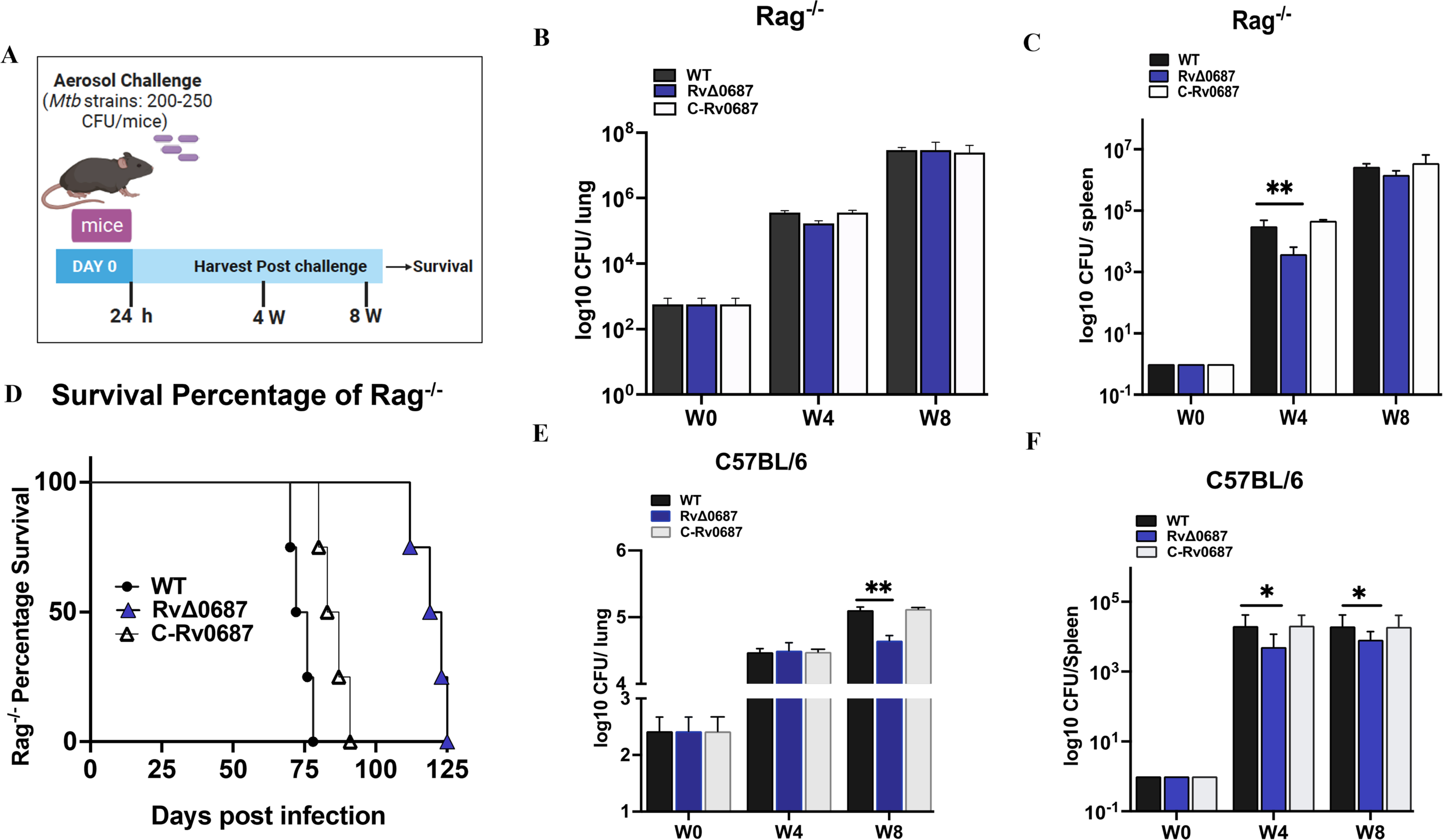
A-F. RvΔ0687 is attenuated for survival in the immunocompromised (Rag^-/-^) and immunocompetent (C57BL/6) Mice model. (A) Infection Strategy. (B-C) Bacterial load in Rag^-/-^ lungs and spleen: The WT, RvΔ0687 and C-Rv0687 strains were used to infect the Rag^-/-^ mice (n= 4 animals per group) and euthanized at W0, W4 and W8. Lungs and spleen were harvested, homogenized, and used to enumerate bacterial load by CFU assay. The experiment was performed with appropriate statistical numbers and significance was calculated using Two-way ANOVA \*\**P<0.01.* (**D) Survival analysis of WT and Rv**Δ**0687.** The survival of mice post infection with WT, C-Rv0687 and RvΔ0687 was performed with n=4 animals per group, and the survival was monitored for 125 days and plotted using the Log rank test. (**E-F) Bacterial load in C57BL/6 lungs and spleen.** The WT, RvΔ0687 and C-Rv0687 strains were used to infect C57BL/6 mice (n= 4 animals per group) and were euthanized at W0, W4 and W8. Lungs and spleen were processed for bacterial enumeration and CFUs were enumerated per lung and spleen. The experiment was performed with appropriate statistical numbers and significance was calculated using Two-way ANOVA (\*\**P<0.01* and \**P<0.1)*.

To further evaluate the role of Rv0687 in *Mtb* pathogenesis *in-vivo*, we infected C57BL/6 mice and euthanized them at 4- and 8-weeks post-infection. Lungs and spleen were harvested, plated and CFUs were enumerated. In lungs though at W4 CFUs were similar for WT, RvΔ0687 and C-Rv0687 strains, differences were observed during late infection at W8, enumerated CFUs for RvΔ0687 were 3-fold lower than WT or C-Rv0687 (**Fig.5E).** Further in spleen compared to WT or C-Rv0687 infected mice at W4 and W8, RvΔ0687 infected mice displayed ∼ 4-fold or 2.5-fold lower CFU’s respectively (**Fig. 5F**).

### 4. Discussion

In this study, we investigated the functional role of *Mtb* Rv0687 that codes for a putative short-chain dehydrogenase. In bacteria including *Mtb*, its reported that short-chain dehydrogenase/reductases (SDRs) aid in managing peroxidase stress, detoxification, *in-vitro*, *in-vivo* survival, drug resistance and overall homeostasis (16,17). Interestingly, in our RNA-seq studies (data not shown) we found that Rv0687 is highly upregulated in *Mtb* treated with NMR711, a compound that we found can inhibit the growth of both drug-sensitive and multidrug-resistant TB (29). Better understanding of role of these SDR’s in *Mtb* pathogenesis will lead to development of drugs and therapeutics in future to combat tuberculosis (TB). To understand the functional role of *Mtb* Rv0687, we created a gene knockout mutant of Rv0687 (RvΔ0687), its complemented strain (C-Rv0687) and characterized the role of Rv0687 in *Mtb* survival and growth under various stress conditions including minimal media, oxidative, nitrosative and drug-mediated stress. We also investigated the function of Rv0687 in *Mtb* infection in BMDMs and mice models. Overall, our study suggests that Rv0687 has a substantial role in *Mtb* pathogenesis in combating oxidative and nitrosative stress, drug tolerance, and host immunity, and is indispensable for pathogenicity of *Mtb* in macrophages and mice models of infection.

Bacterial growth and survival are influenced by the nature and amount of nutrients available in the growth media (35).While performing growth kinetics we observed that in nutrient-rich conditions, the growth of RvΔ0687 was indistinguishable from that of WT and C-Rv0687 *in-vitro*. However, when grown in dextrose-containing minimal media supplemented with glycerol, polysorbate and 0.2% dextrose, we found that WT, RvΔ0687 and C-Rv0687 strains were able to survive normally till day 8 and declined gradually until day 12. This suggests that the addition of the dextrose, which offers a limited nutrient source, could sustain bacterial growth for an initial period, and the bacterial strains might have efficiently utilized the available dextrose for survival during early time points as described previously (36–38). However, the dextrose present in the media was insufficient to support bacterial growth beyond day 8, which was evidenced by the inability of any of the *Mtb* strains to grow after 8 days until 12 days post-inoculation. Our findings parrels with observations made with the gene knockout mutant of *rel_Mtb_*, which displayed a significantly lower growth rate than the WT in minimal media. As shown in previous studies *Mtb* ReI_Mtb_ protein is required for long-term survival of pathogenic mycobacteria under starvation conditions (39). These results suggest that the carbon source used in the growth media might alter the growth rate, which can impact *Mtb* survival for a prolonged time. Taken together, our findings highlight the contribution of the Rv0687 gene in the adaptation of *Mtb* to grow under nutrient-limited conditions.

Since tolerance to ROS and RNS play essential roles in bacterial pathogenesis, we further investigated the response of *Mtb* strains against stress-generating agents (40). In our pilot experiment (data not shown) we used 7H9 complete media supplemented with glycerol, polysorbate and OADC to perform the stress response studies, and we did not observe significant growth defect between the WT, RvΔ0687 and C-Rv0687 strains treated with 5mM H_2_O_2_. Therefore, we tested the phenotype of the *Mtb* strains in 7H9 dextrose minimal media and found that the RvΔ0687 was more sensitive to ROS-generating agents, as evidenced by a decline in its growth compared to WT and C-Rv0687 (41). Our results showed that RvΔ0687 is sensitive to ROS (H_2_O_2_ and tBOOH). Our results are further supported by previous microarray studies that shows Rv0687 is upregulated on *Mtb* treatment with H2O2 (42). Like *Mtb sigJ, mel2* and *sod* that are known to play role in providing resistance against ROS and *Mtb* pathogenesis (41,43,44), Rv0687 might is also helps *Mtb* in coping up the ROS and is required for *Mtb* pathogenesis.

In addition, we investigated the response of RvΔ0687 to RNS by treating bacterial cultures with SNAP and SN. The increased sensitivity of RvΔ0687 to SNAP treatment suggests that the Rv0687 gene may play a role in *Mtb* defense against nitrosative stress induced by SNAP, which is a nitric oxide donor, as has been shown for the gene knockout mutant of Rv2617c of *Mtb* involved in virulence (45). Our findings also suggest that RvΔ0687 can tolerate exposure to SN up to 10mM concentrations; this was like the findings of a study by Suwarna et al., which showed that *Mtb* can resist and survive in the presence of a nitrite concentration up to 10mM in broth media (46). So might be higher concentration of sodium nitrite is required to generate sufficient RNs species to inhibit the growth of *Mtb*. Additionally, we tested the response of RvΔ0687 to exposure to transitional metals such as copper, zinc, and iron, which can induce oxidation reactions. However, we could not find a significant difference in the survival between the mutant and WT or complemented strains.

The *Mtb* being an intracellular pathogen grows inside the phagosomes of macrophages where oxygen and nitrogen intermediates are produced; having such evasion mechanisms would enable *Mtb* to survive under stress conditions. SDRs in *Mtb* are involved in regulating peroxidase stress and detoxification. Interestingly, a homolog of Rv0687 in other mycobacteria is characterized as a mycofactocin-associated dehydrogenase with non-exchangeable NAD-cofactors. Homolog of Rv0687 in *M. paratuberculosis* is a putative carveol dehydrogenase and its molecular structure has been reported. Accordingly, carveol has been shown to have an antioxidant role (47). This suggests that Rv0687 might also be involved in redox reactions or synthesis of cellular metabolites that can play an important role in the detoxification of ROS or RNS in *Mtb* and can be further explored in the future. Therefore, understanding the specific mechanisms by which the Rv0687 gene confers protection against ROS and RNS could provide valuable insights into the bacterial stress response relating to the pathogenesis of *Mtb*.

We also investigated the impact of Rv0687 on the redox homeostasis of *Mtb*. In general, redox homeostasis, involving the balance between reduced NADH and oxidized NAD^+^, plays a key role in maintaining cellular redox equilibrium, which is crucial for bacterial survival and adaptation to various environmental stresses, including oxidative and nitrosative stresses. (32,48–50). Hence, any disruption in redox homeostasis can have a significant impact on bacterial physiology and survival (51,52). We found that the NADH and NAD^+^ concentrations, as well as the NADH/NAD^+^ ratio, were comparable between RvΔ0687 and the WT and C-Rv0687 strains. These results were consistent with the phenotype of the DosR regulon mutant of *Mtb* known to assist *Mtb* in metabolic homeostasis and recovery from dormancy but DosR mutant of *Mtb* exhibited similar NADH and NAD^+^ levels as the WT during aerobic growth conditions at early time points. There is a possibility of plasticity in SDRs that can compensate for the function of Rv0687, as it has been shown that deletion of a single succinate dehydrogenase (ndhA) gene, which is an SDR, doesn’t have major redox potential in *Mtb* (32,51). These observations suggest that Rv0687 might not be directly involved in regulating the balance between NADH and NAD^+^ in *Mtb* under the conditions tested. It is possible that redundant or alternative pathways are activated to ensure the stability of the NADH/ NAD^+^ ratio in the RvΔ0687 strain or might be required under certain specific unknown conditions. Further research is warranted to elucidate the precise functions and interactions of Rv0687 with other redox-related components and regulatory networks in *Mtb*.

As SDR Rv0148 of *Mtb* has been associated with the sensitivity to antibiotics, in the current study, we assessed the sensitivity of WT, RvΔ0687 and C-Rv0687 strains against anti-*Mtb* antibiotics, using alamar blue assay. The results demonstrated distinct drug susceptibility patterns for the RvΔ0687 mutant. Notably, compared to the WT and C-Rv0687 strains, the RvΔ0687 showed increased sensitivity to two drugs: NMR711 and delamanid. Conversely, the RvΔ0687 strain exhibited no significantly different sensitivity than the WT and C-Rv0687 strains against INH, BDA, and RIF. This indicates that the Rv0687 plays a role in bacterial sensitivity, particularly to DEL and NMR711 (53). It correlates with our unpublished data that Rv0687 was upregulated in *Mtb* samples treated with NMR711. There is possibility that it might work synergistically with delamanid, recently approved TB drug but further studies are required. Further studies are also needed to elucidate the underlying mechanisms by which Rv0687 influences the bacterial susceptibility to these drugs.

During *Mtb* infection, recognition of pathogen by macrophages induces changes in the expression of cytokines that impact the elimination or progression of infection (54). To assess the infection and *in-vivo* survival of WT, RvΔ0687 and C-Rv0687 strains, we used a mouse BMDMs infection model and monitored the intracellular bacterial survival after infection. We observed that RvΔ0687 was defective for intracellular survival compared to WT and C-Rv0687 strains in BMDMs. The current results were consistent with the findings in another SDRs and Rv2159c alkylhydroperoxidase of *Mtb*. The Rv0148 and Rv2159 mutants were defective for intracellular survival, although it induced pro-inflammatory cytokines in a macrophage model (15,16). Compared to uninfected, we have observed significant changes in the production of ROS and RNS by infected BMDMs though there were no significant changes among WT vs RvΔ0687 strains.

Further, *Mtb*-infected phagocytes generate ROS and RNS, which influence the activation of T cell responses, production of pro-inflammatory cytokines, induce autophagy, and apoptosis (10). We analyzed the levels of IL-1β, TNF-α, IP-10 and MIP-1α cytokines in the supernatants of BMDMs infected with *Mtb* strains. Strikingly, we observed that RvΔ0687-infected BMDMs showed significantly higher TNF-α and MIP-1α levels compared to both WT and C-Rv0687 strains. MIP-1α/CCL3 is a chemotactic chemokine secreted by macrophages to recruit inflammatory cells and maintain effector immune response (55). This suggests that the Rv0687 gene might influence the host immune response during *Mtb* infection in macrophages and in mice. To investigate the *in-vivo* survival and virulence of RvΔ0687, in comparison to the WT, and C-Rv0687 strains, we used two mice models of pulmonary infection. In the Rag^-/-^ mice, which lack functional T and B cells, all the tested *Mtb* strains exhibited similar bacterial burden in the lungs at 4 weeks. However, in the spleen, RvΔ0687 displayed a significantly lower bacterial growth compared to WT and C-Rv0687 at this time. This suggests that Rv0687 may play a role in the dissemination of *Mtb* primary infection from lung to spleen and early establishment of *Mtb* infection in the spleen of immunocompromised mice. But at 8 weeks post-infection, all the strains showed a comparable bacterial burden in both the lungs and spleen, indicating that the RvΔ0687 mutant was able to overcome the initial difficulty in spleen colonization. In addition, RvΔ0687 mutant-infected Rag^-/-^ mice survived for a longer duration than the WT-infected mice, suggesting that Rv0687 have some role in the *in-vivo* survival of *Mtb* in the absence of functional T and B cells. Interestingly, in C57BL/6 mice, a similar bacterial load was observed in the lungs of WT, RvΔ0687 and C-Rv0687 strains after 4 weeks of infection. However, at 8 weeks post-infection, RvΔ0687 displayed a lower bacterial burden in the lungs and reduced bacterial load in the spleen at both 4- and 8-weeks post-infection. This suggests that the Rv0687 gene may have significant role in counteracting host adaptive immunity and in the *in-vivo* survival and virulence of *Mtb* in immunocompetent mice. Together, these findings shed light on the complex interplay between *Mtb* and the host immune system, emphasizing the significance of the Rv0687 gene in modulating *Mtb* survival in mouse lungs and spleen.

### Future Studies

Mtb lives in diverse environments within the host macrophages and is constantly exposed to stressful conditions imposed by the host. To survive and replicate inside the macrophage *Mtb* has acquired various evasion mechanisms including to combat host-generated oxidative and reductive stress (ROS, RNS etc). Further studies involving biochemical characterization of the SDRs reported in this study and its interaction with other regulatory networks will shed light on its role in bacterial adaptation to the host environment, tolerance to antibiotics and in *Mtb* pathogenesis.

**Table 1:** Minimal Inhibitory Concentration (MIC) for WT, RvΔ0687 and C-Rv0687 strains: Isoniazid. (INH) 1-0.007 μg/ml, Delamanid (DEL) 1-0.007 μg/ml, NMR711 (50-0.39 μg/ml, Bedaquiline (BDA)-2-0.015 μg/ml, Rifampicin (RIF)- 8-0.062 μg/ml. The above-mentioned concentrations were tested and reported the MIC based on the color change with alamar blue assay.

| Strain Name | INH<br>(1-0.007<br>μg/ml) | DEL<br>(1-0.007<br>μg/ml) | NMR711<br>(50-0.39<br>μg/ml) | BDA<br>(2-0.015<br>μg/ml) | RIF<br>(8-0.062<br>μg/ml) |
| --- | --- | --- | --- | --- | --- |
| WT | >1 | >1 | >50 | >2 | >8 |
| **RvΔ0687** | >1 | ≥ 0.062 | ≥ 0.390 | >2 | >8 |
| **C-Rv0687** | >1 | >1 | >50 | >2 | >8 |

## Data availability statement

The supporting data of this study are available in the supplementary material of this article

## Ethics statement

The animal protocol was approved by the institutional IACUC committee of University of Texas at El Paso under the protocol (A-202004-3). All the animals were maintained according to the guidelines of IACUC.

## Author Contributions

ST has initiated the study. GB and ST designed the experiments; GB, MKM, DK performed the experiments. GB and ST conceptualized the data and GB plotted graphs. GB and ST drafted the manuscript. GB, ST, DK and MKM edited, revised, and approved the final version of manuscript.

## Supplementary Figures

**Figure S1. Construction of gene deletion mutant of Rv0687 and its complementation:**

Rv0687 was deleted in *Mtb* using specialized transduction and confirmed by three primer PCR A-C. (A) Three primer PCR product confirmation (B) Gene specific PCR for *Mtb* Rv0687 gene.

**(C)** PCR for Kanamycin marker to confirm the complemented strain. This was performed using H37Rv, CDC1551, RvΔ0687 and C-Rv0687 strains.

**Figure S2. Effect of Sodium Nitrite Stress on *Mtb* and** RvΔ068: The WT, RvΔ0687 and C-Rv0687 strains were grown in 7H9 dextrose media and exposed to 2 or 10mM sodium nitrite and all the bacterial strains were monitored for growth at 0, 24, 48, 72 and 96h and CFUs were estimated after 3 weeks of incubation and calculated for per ml of bacterial culture.

**Figure S3: Response of WT, Rv**Δ**0687 and C-Rv0687 to Metal Stress:** The WT, RvΔ0687 and C-Rv0687 strains were grown in 7H9 dextrose media and exposed to 75μM CuSO_4,_ 75μM ZnSO_4_ or 0.05% FeSO_4_ and the cultures were monitored for growth at 0, 24 and 72h. The CFUs were analyzed by serially diluting the cultures and plating on 7H10 plates and CFUs were calculated per ml of bacterial culture.

**Figure. S3. MIC using Alamar Blue to test drug-sensitivity:** Anti-TB drugs **(A)** NMR711, **(B)** Delamanid, **(C)** Isoniazid, **(D)** Rifampicin and **(E)** Bedaquiline drugs were added onto top row wells, and two-fold serial dilutions were performed with 7H9 complete media broth till it reaches to least concentrations. A 0.1 OD of 100 µl bacterial culture and 1x alamar blue were added to each well. Blue color represents the reference wells (Blank), pink color represents the concentration of drug in which bacteria is alive and bluish pink color represents the MIC of bacteria where it tends to inhibit. The picture show assay plates at Day 7 post incubation.

**Figure. S4 A-B. RNS and ROS production by BMDM in response to infection**

**(A) Assessing the RNS production by BMDMs during Infection:** The RNS production was measured in the cell free supernatants of BMDM infected with WT, RvΔ0687 or C-Rv0687 strains at Day 0, 1, 5 and 7 (**B) Assessing the ROS production by BMDMs during Infection:** The production of ROS was measured at Day 0, 1, 3 and 7 in the post infected lysates of BMDM infected with WT, RvΔ0687 or C-Rv0687 strains using optical density readings, using uninfected lysates as a control.

**Table S1:**
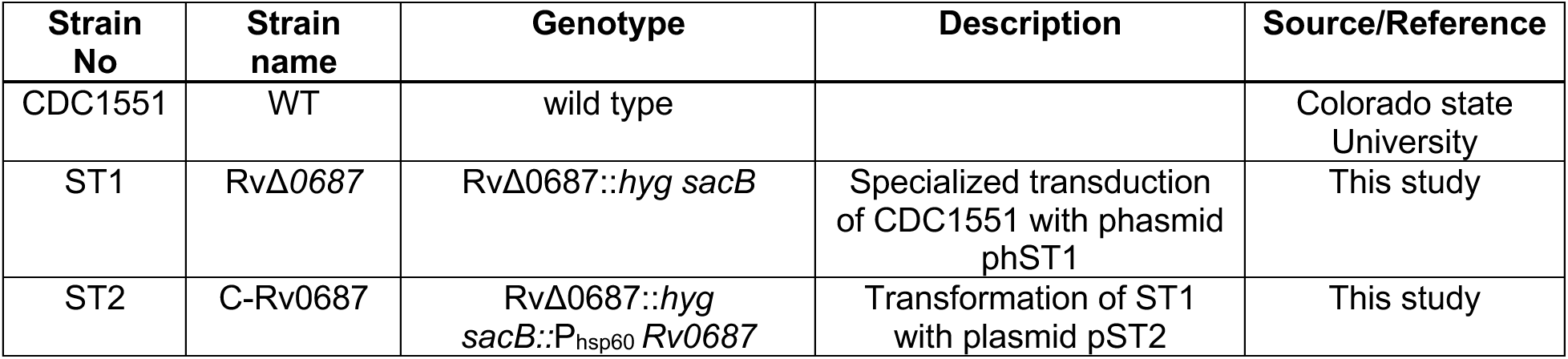
List of strains used in the current study. Abbreviations used *hyg*: hygromycin casette, *sacB*: counter selectable marker.

| Strain No | Strain name | Genotype | Description | Source/Reference |
| --- | --- | --- | --- | --- |
| CDC1551 | WT | wild type |  | Colorado state University |
| ST1 | RvΔ0687 | RvΔ0687::*hyg sacB* | Specialized transduction of CDC1551 with phasmid phST1 | This study |
| ST2 | C-Rv0687 | RvΔ0687::*hyg sacB::P<sub>hsp60</sub> Rv0687* | Transformation of ST1 with plasmid pST2 | This study |

**Table S2:** List of plasmids and phasmids used in the current study: Abbreviations used *hyg*^R^: hygromycin resistance, *suc*^S^: Sucrose sensitive, *kan*^R:^ Kanamycin resistance.

| Plasmids/phages | Description | Reference |
| --- | --- | --- |
| P004s | Suicide recombination delivery vector carrying *hyg<sup>R</sup> sacB* for gene disruption, *hyg<sup>R</sup>* | Jain et., al 2004 |
| pST1 | P004s containing homologous left and right arm to delete Rv0687, *hyg<sup>R</sup>, suc<sup>S</sup>* | This study |
| phAE159 | Conditionally replicating shuttle phasmid vector | Jain et., al 2004 |
| phST1 | Phsmid with flanking regions to delete Rv0687 by homologous recombination, *hyg<sup>R</sup>, suc<sup>S</sup>* | This study |
| pMV361 | Integrative mycobacterial shuttle vector with hsp60 promoter carrying *kan<sup>R</sup>* resistant gene | Strover et., al 1991 |
| pST2 | Rv0687 was cloned in pMV361, *kan<sup>R</sup>* | This study |

**Table S3:**
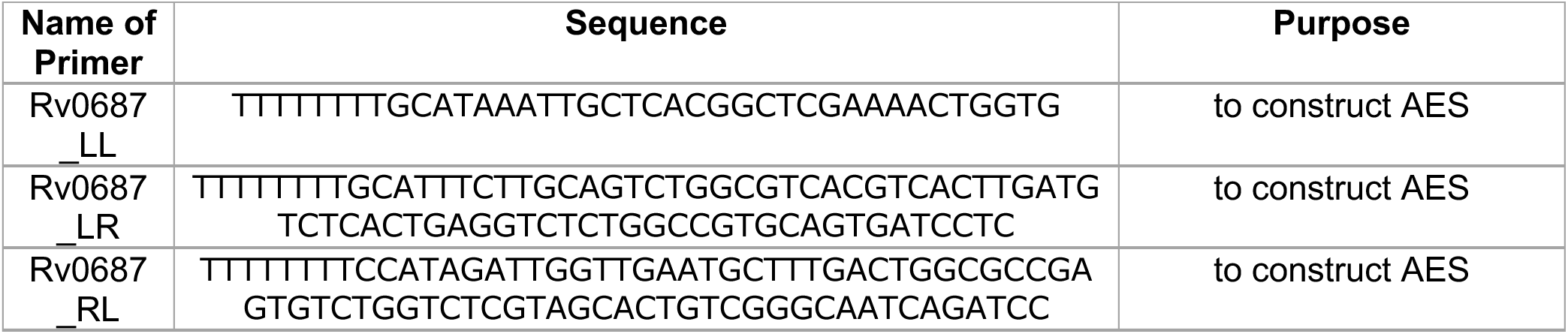
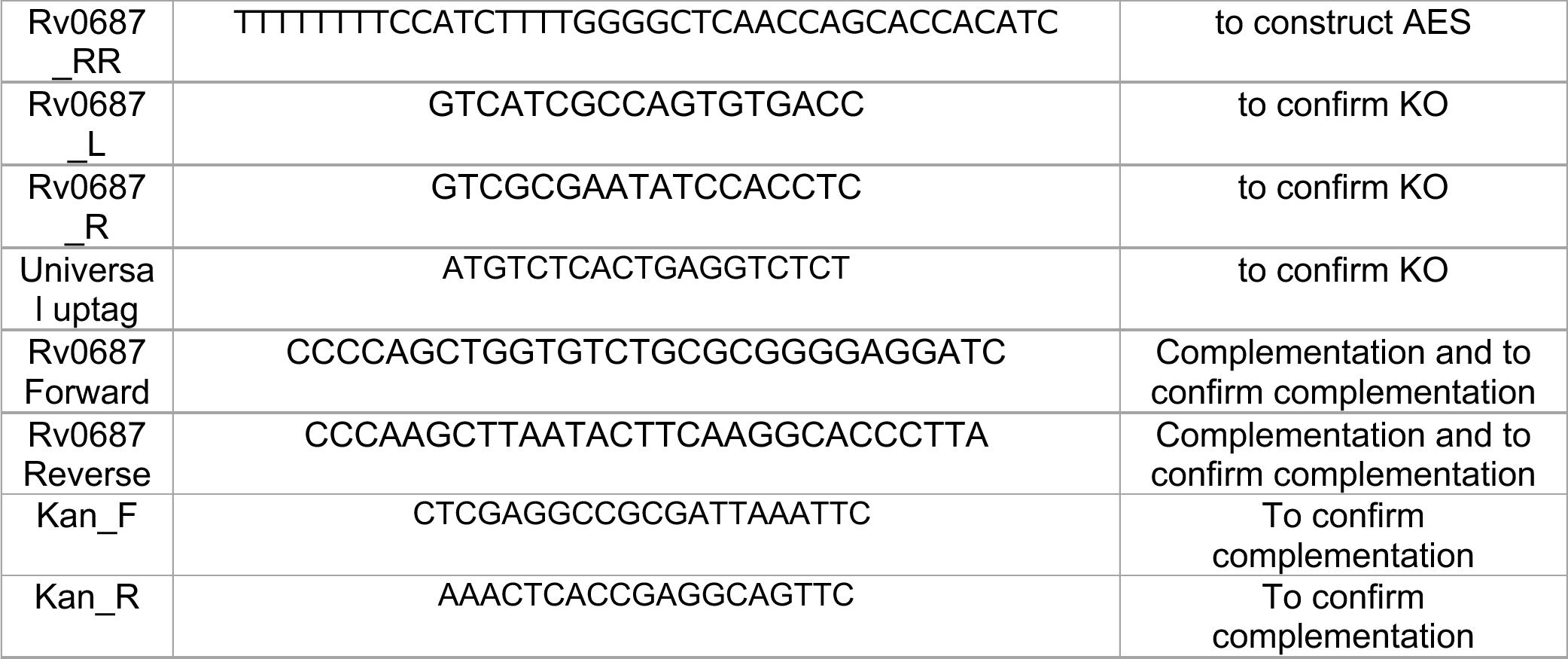
Primers used in the current study. Abbreviations used AES: Allelic exchange substrate KO-knockout deletion strain.

